# Mini viral RNAs act as innate immune agonists during influenza virus infection

**DOI:** 10.1101/385716

**Authors:** Aartjan J.W. te Velthuis, Joshua C. Long, David L.V. Bauer, Rebecca L.Y. Fan, Hui-Ling Yen, Jane Sharps, Jurre Y. Siegers, Marian J. Killip, Hollie French, Maria José Oliva-Martín, Richard E. Randall, Emmie de Wit, Debby van Riel, Leo L.M. Poon, Ervin Fodor

**Author notes:** These authors contributed equally to this work (Aartjan J.W. te Velthuis, Joshua C. Long, David L.V. Bauer).

## Abstract

Influenza A virus infection usually causes a mild to moderately severe respiratory disease in humans. However, infection with the 1918 H1N1 pandemic or highly pathogenic avian influenza viruses (HPAIV) of the H5N1 subtype, can lead to viral pneumonia, systemic disease and death. The molecular processes that determine the outcome of influenza virus infection are multifactorial and involve a complex interplay between host, viral, and bacterial factors^1^. However, it is generally accepted that a strong innate immune dysregulation known as ‘cytokine storm’ contributes to the pathology of pandemic and avian influenza virus infections^2–4^. The RNA sensor Retinoic acid-inducible gene I (RIG-I) plays an important role in sensing viral infection and initiating a signalling cascade that leads to interferon (IFN) expression^5^. Here we show that short aberrant RNAs (mini viral RNAs; mvRNAs), produced by the viral RNA polymerase during the replication of the viral RNA genome, bind and activate the intracellular pathogen sensor RIG-I, and lead to the expression of interferon-β. We find that erroneous polymerase activity, dysregulation of viral RNA replication, or the presence of avian-specific amino acids underlie mvRNA generation and cytokine expression in mammalian cells and propose an intramolecular copy-choice mechanism for mvRNA generation. By deep-sequencing RNA samples from lungs of ferrets infected with influenza viruses we show that mvRNAs are generated during infection of animal models. We propose that mvRNAs act as main agonists of RIG-I during influenza virus infection and the ability of influenza virus strains to generate mvRNAs should be considered when assessing their virulence potential.

---

The negative sense viral RNA (vRNA) genome segments of influenza A viruses, as well as the complementary RNA (cRNA) replicative intermediates, contain 5′ triphosphates and partially complementary 5′ and 3′ termini that serve as the viral promoter for replication and transcription of the viral RNA genome^6^. RIG-I has been shown to bind and be activated by the dsRNA structure formed by the termini of influenza virus RNAs^7,8^. However, it remains unclear how RIG-I gains access to this dsRNA structure. Both vRNA and cRNA are assembled into ribonucleoprotein complexes (vRNP and cRNP, respectively) in which the viral RNA polymerase, a heterotrimeric complex of the viral proteins PB1, PB2 and PA, associates with the partially complementary termini, while the rest of the RNA is bound by oligomeric nucleoprotein (NP)^6^ (Fig. 1a). The tight binding of the 5′ and 3′ termini of vRNA and cRNA by the RNA polymerase^9^ is likely to preclude an interaction with RIG-I. Moreover, it has been demonstrated that IFN expression is triggered only in a fraction of influenza virus infected cells^10,11^, suggesting that influenza viruses efficiently hide their genome segments during infection by replicating them in the context of RNPs^11^. This led to the proposal that an aberrant RNA replication product might be binding to RIG-I and triggering IFN expression^12^. Indeed, the influenza virus polymerase is known to generate defective interfering (DI) RNAs, which are ≥178 nt long subgenomic RNAs generated during high multiplicity infections^13^, and small viral RNAs (svRNAs), which are 22-27 nt long and correspond to the 5′ end of vRNA segments. However, svRNAs have been shown not to be involved in the induction of antiviral cellular defences^14^ and DI RNAs assemble into RNP structures (Fig. 1a), as demonstrated for a 248 nt long DI RNA^15^, potentially precluding their interaction with RIG-I. Therefore, it remains unclear what kind of viral RNA species is recognised by RIG-I (Fig. 1A) and why different influenza virus strains trigger dramatically different levels of IFN expression^2,3,16^.

**Figure 1.**
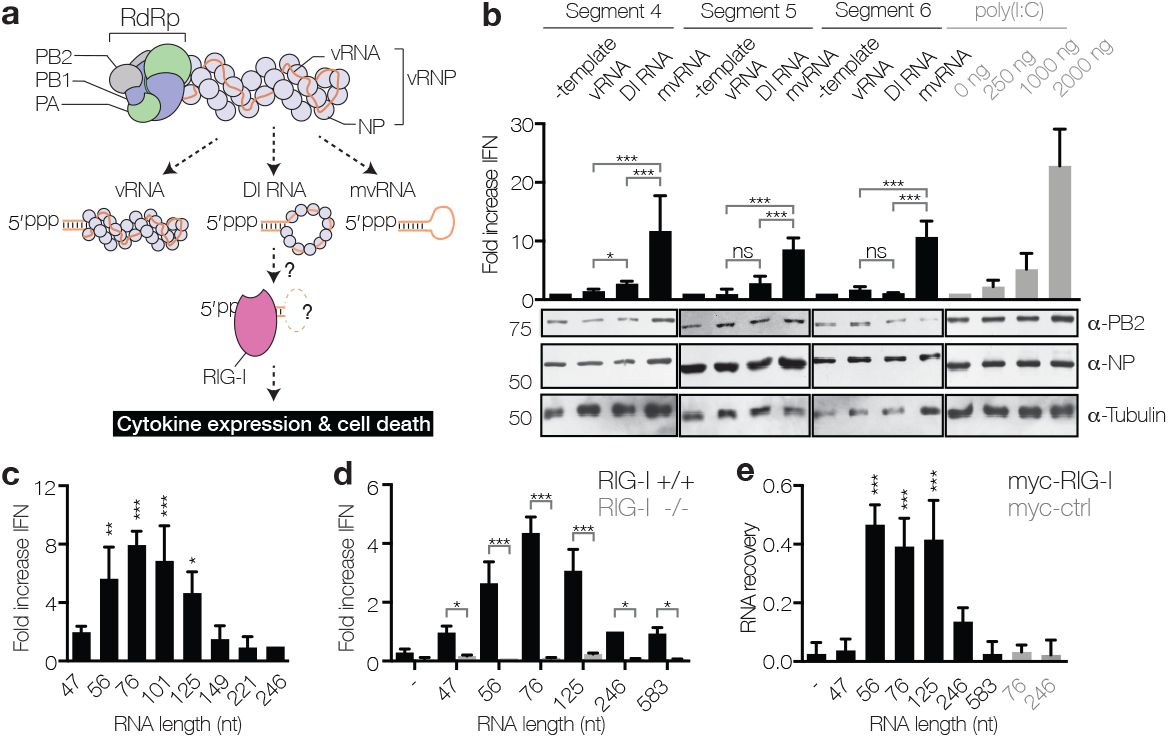
mvRNAs of influenza A virus are bound by RIG-I and induce IFN expression. (**a**) Models of the influenza virus ribonucleoprotein (vRNP) complex and potential activators of RIG-I. (**b**) Analysis of IFN-β promoter activity induced by the replication of segment 4, 5 or 6 vRNAs, DI RNAs or mvRNAs or by the transfection of poly(I:C). (**c**) IFN-β promoter activity induced by the replication of engineered, segment 5-based, short RNAs in HEK 293T cells or (**d**) in wild-type or HEK 293 RIG-I knockout cells. (**e**) Binding of segment 5-based RNAs to myc-tagged RIG-I or mouse EGF control protein (myc-ctrl). In all panels, error bars indicate standard deviation with n=3. P-values were determined using ANOVA, are shown compared to 246 nt RNA in Fig. 1C and myc-ctrl with 246 nt mvRNA in Fig.1E, and are indicated as * P ≤ 0.05, ** P ≤ 0.01, *** P ≤ 0.001.

Engineered viral RNAs shorter than 149 nt but containing both the 5′ and 3′ termini of vRNAs can be transcribed and replicated in cells by the viral polymerase in the absence of NP17, suggesting that they do not form canonical RNP structures. We call these short viral RNAs mvRNAs (Fig. 1a). To investigate which class of viral RNA is responsible for triggering IFN expression, we expressed a full-length segment 4 (hemagglutinin encoding) vRNA (1775 nt long) or its truncated versions, a 245 nt long DI RNA and 77 nt long mvRNA, in HEK 293T cells together with viral polymerase, NP and a luciferase reporter to measure the activation of the IFN-β promoter (Fig. 1b). We found that the expression of mvRNAs induced significantly higher IFN expression than full-length vRNA or DI RNA, comparable to the levels induced by transfection of 2 μg of poly(I:C), a known activator of IFN expression^18^. Similar results were obtained with segment 5 and 6 (NP and neuraminidase encoding, respectively) vRNAs and their truncated DI RNA and mvRNA versions (Fig. 1b). To determine the optimal mvRNA length that triggers activation of the IFN-β promoter, we expressed 47 to 246 nt long vRNAs derived from segment 5 together with viral polymerase and NP and measured the activity of the IFN-β promoter. We found that the replication of 56 to 125 nt long mvRNAs resulted in significantly higher IFN-β promoter activity than the replication of RNAs shorter than 56 nt or longer than 125 nt (Fig. 1c, Fig. S1a and b).

To address whether these engineered short mvRNAs triggered IFN expression via RIG-I, we co-expressed viral RNAs with polymerase and NP in HEK 293T RIG-I knockout (RIG-I -/-) or control (RIG-I +/+) cells engineered to express luciferase in response to the activation of the IFN-β promoter. We found that 56 to 125-nt long mvRNAs induced only background levels of luciferase in RIG-I -/- cells, even though the transfection of a plasmid expressing RIG-I or 2 μg of poly(I:C) resulted in significant activation of the IFN-β promoter (Fig. 1d, Fig. S1c). By contrast, significant levels of luciferase activity were detected in RIG-I +/+ cells (Fig. 1d). mvRNAs of 56 to 125 nt induced the strongest activation of the IFN-β promoter, in agreement with the data above (Fig. 1c and d, Fig. S1c and d). To address whether mvRNAs trigger the activation of IFN-β expression through binding to RIG-I, we immunoprecipitated myc-RIG-I from cells expressing RNAs of 47 to 583 nt and analysed the amounts of co-immunoprecipiated RNAs. We observed that mvRNAs of 56 to 125 nt were specifically enriched in RIG-I immunoprecipitates (Fig. 1e, Fig. S1e). No mvRNAs were detected in mouse myc-EGF immunoprecipitates used as a negative control (Fig. 1e). To test if mvRNAs also activate RIG-I, we incubated purified myc-RIG-I (Fig. S1f) with an *in vitro* transcribed and gel purified 76 nt mvRNA and measured ^32^P_i_ release in an ATPase assay. We found that a triphosphorylated 76 nt mvRNA induced higher levels of ATPase activity than a dephosphorylated 76 nt mvRNA, while no ATPase activity was observed when we incubated a RIG-I mutant with the triphosphorylated 76 nt mvRNA (Fig. S1g). Overall, these results demonstrate that mvRNAs longer than 47 and shorter than 125 nt are bound by RIG-I, which results in RIG-I activation and the induction of IFN-β expression. This is in agreement with findings that reconstitution of full-length influenza virus vRNPs leads to only low levels of IFN expression unless the cells are pre-treated with IFN^19^ and the hypothesis that aberrant replication products trigger the IFN induction cascade^12^.

We next asked whether mvRNAs are made during influenza virus infection. We infected HEK 293T cells with influenza A/WSN/33 (H1N1) (abbreviated as WSN) and analysed viral RNAs by RT-PCR of segment 1 (encoding PB2), RT-PCR of all segments using universal primers (Fig. S2a), or deep-sequencing of the total small RNA fraction (RNAs 17 to 200 nt in length) (Fig. S2b). We found only very low levels of mvRNAs and, consistently, observed no significant IFN expression (Fig. 2a and b). We hypothesised that mvRNAs are only generated as a consequence of dysregulated viral RNA replication. To test this, we overexpressed viral RNA polymerase from plasmids prior to infection to generate an imbalance between polymerase and NP levels, which is known to induce innate immune signalling^20^. As shown in Fig. 2a and b, under this condition we found significantly higher levels of mvRNAs and IFN expression, while simultaneous overexpression of NP and polymerase reduced mvRNA and IFN production (Fig. 2a). We verified the identity of mvRNAs using gel isolation and Sanger sequencing (Fig. S2c) as well as deep sequencing (Fig. 2b). We found that the majority of mvRNAs were derived from the PB1-, HA-, NP- and NA-encoding vRNA segments (Fig. 2c) and that mvRNAs had a size distribution with a peak around 55 to 64 nt (Fig. 2d, Fig. S2d). In addition to mvRNAs, we also identified complementary mini viral RNAs (mcRNAs).

**Figure 2.**
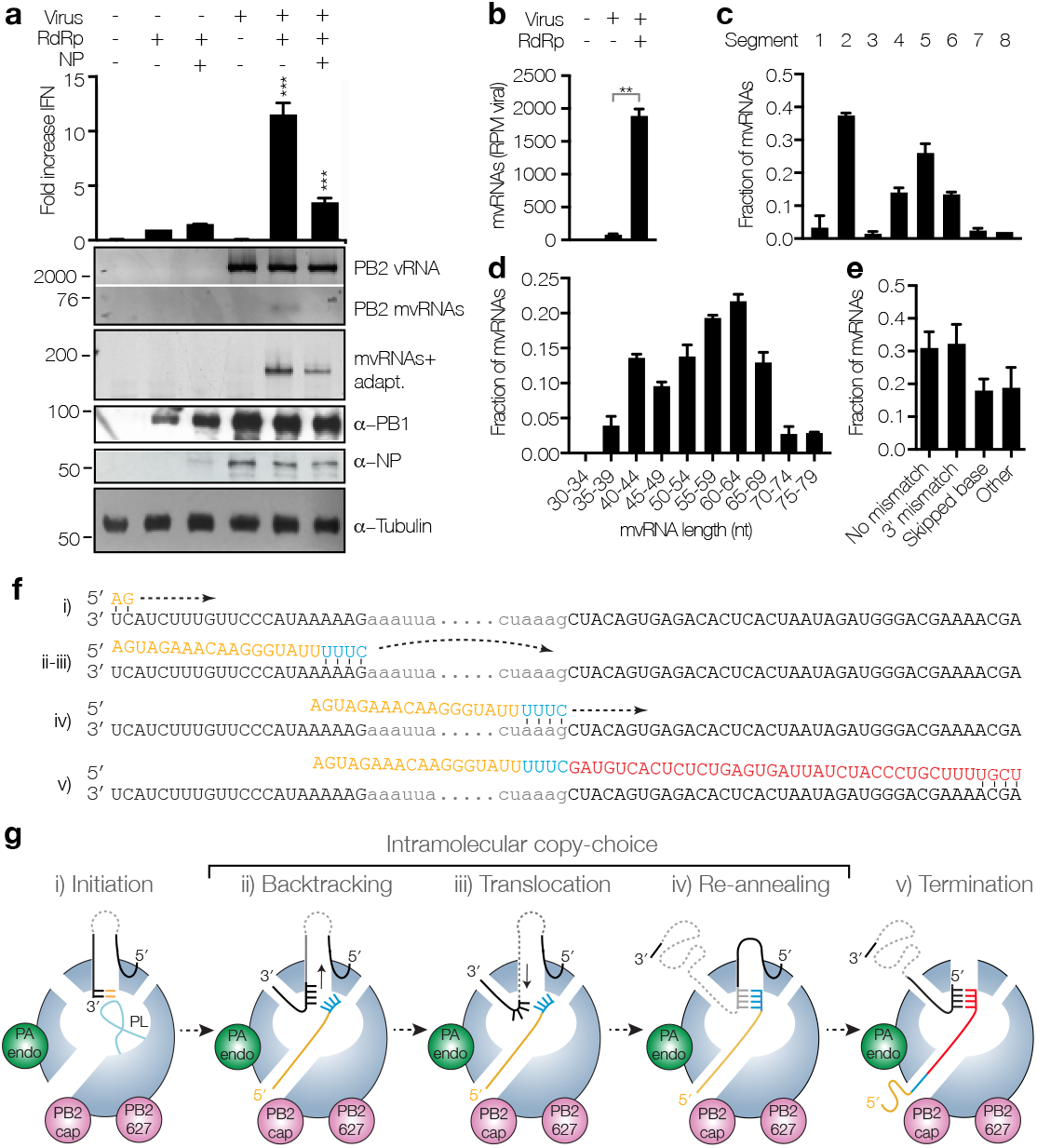
Dysregulation of RNA replication in cells infected with WSN results in the generation of mvRNAs. (**a**) Analysis of IFN-β promoter activity and steady state vRNA and mvRNA levels in WSN infections following overexpression of viral polymerase or viral polymerase and NP. mvRNAs were also amplified with universal primers containing adapters for sequencing (mvRNAs+adapt). (**b**) Quantitation of mvRNAs using deep sequencing, expressed as reads per million (RPM). (**c**) mvRNA distribution per genome segment. (**d**) Size distribution of mvRNAs. (**e**) mvRNA distribution per type of intramolecular copy-choice mechanism. (**f**) Example of mvRNA formation through an intramolecular copy-choice mechanism involving a 3′ mismatch. (**g**) Model of mvRNA formation by the polymerase (model adapted from^6^). In all panels, error bars indicate standard deviation with n=3. P-values were determined using ANOVA and shown compared to lane 3 in Fig. 2A and using an unpaired t-test in Fig. 2b, and are indicated as in Fig. 1.

Analysis of mvRNA sequences suggests that mvRNAs are generated via an intramolecular copy-choice mechanism that tolerates 3' mismatches or skipped bases (Fig. 2e and f). As shown in Fig. 2g, the generation of mvRNAs can be explained by the separation of the template and nascent product RNAs by backtracking^21^, followed by template translocation until base pairing between template and nascent product RNA is re-established (Fig. 2f and g). This process may be induced by an imbalance between viral polymerase and NP levels (Fig. 2a, Fig. S3a). In light of these results, it is important to note that the existence of mvRNAs has likely been overlooked till now, because i) RNA isolation protocols vary in their capacity to recover small RNAs, ii) RT-PCR products from mvRNAs form a diffuse fast-migrating band on standard agarose gels that may be mistaken for primer-dimers, iii) conventional RNA deep sequencing protocols discard short library fragments, and iv) standard ligation-based deep sequencing protocols for small RNAs require a 5′ monophosphate and do not detect viral transcripts with a 5′-triphosphate group.

In humans, infection with the 1918 H1N1 pandemic virus or H5N1 HPAIV lead to strong innate immune activation^2,3,16^. To address whether mvRNAs could contribute to this phenomenon, we investigated the replication of a 246 nt RNA by the polymerase of these viruses. We found that the polymerases of the highly virulent A/Brevig Mission/1/18 (H1N1) (abbreviated as BM18) pandemic virus and the A/duck/Fujian/01/02 (H5N1) (abbreviated as FJ02) HPAIV generated higher levels of mvRNAs than the polymerases of WSN and A/Northern Territory/60/68 (H3N2) (abbreviated as NT60) viruses, even in the presence of high NP concentrations (Fig. 3a). No mvRNAs were observed in a control with an inactive WSN polymerase that had two point mutations in the polymerase active site (PB1a). Using gel isolation and Sanger sequencing we confirmed that the mvRNAs produced by the BM18 polymerase were similar to the WSN mvRNAs (Fig. S3b). Isolation of total RNA from cells expressing polymerase of the BM18 or FJ02 virus and its subsequent transfection into HEK 293T cells resulted in significantly higher IFN-β promoter activity compared to when RNA from cells expressing WSN, NT60, or active site mutant WSN PB1a polymerase was transfected (Fig. 3a).

**Figure 3.**
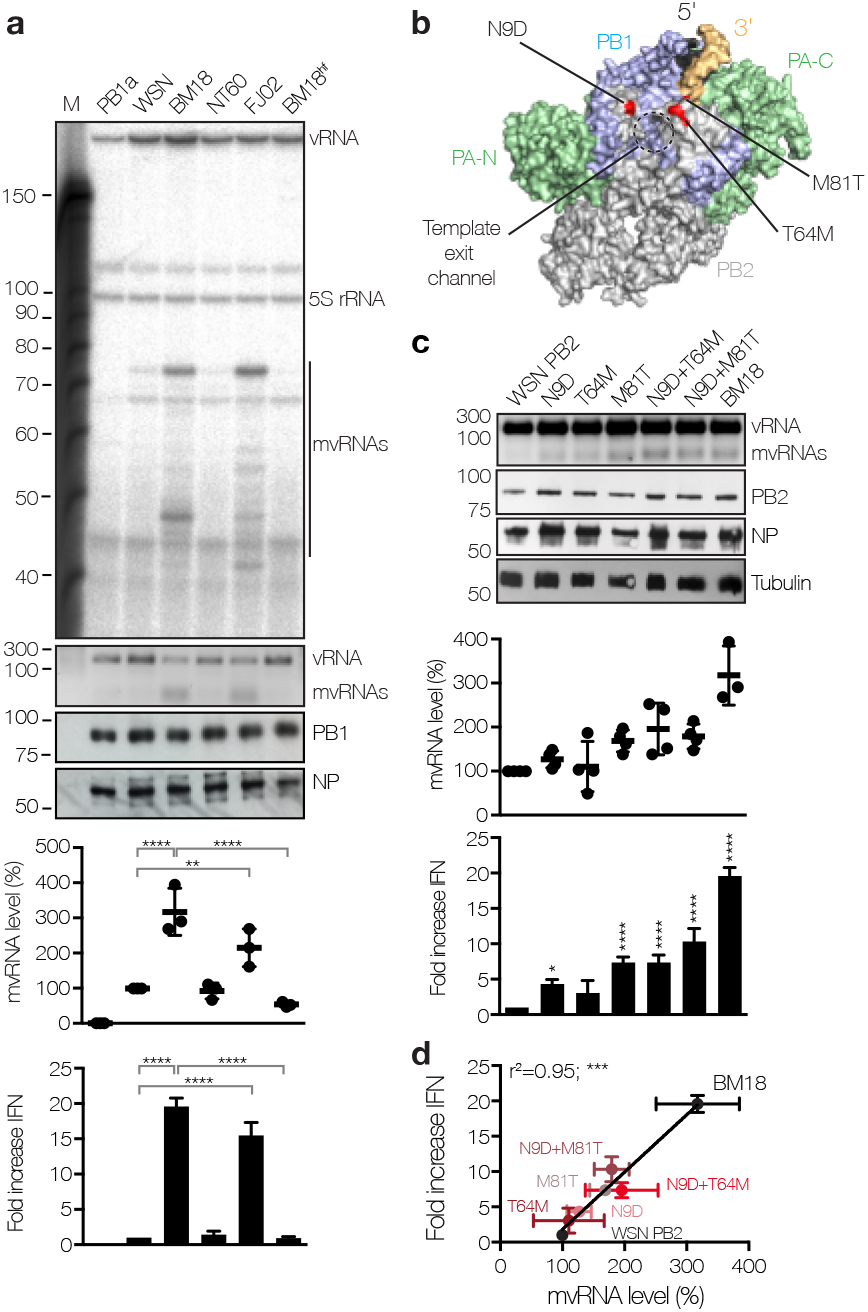
The PB2 polymerase subunit of highly virulent influenza A viruses promotes mvRNA synthesis. (**a**) Analysis of mvRNA levels during the replication of a segment 5-based 246 nt RNA template by the WSN, BM18, NT60 and FJ02 polymerases, and IFN-β promoter activity induced by the transfection of total RNA isolated from these cells into reporter HEK 293T cells expressing luciferase. (**b**) Location of PB2 amino acid residues 9, 64 and 81 in the bat influenza A virus polymerase structure (PDB 4WSB). (**c**) Analysis of the effect of PB2 mutations on mvRNA formation and IFN-β promoter activity induced after transfection of total RNA isolated from these cells into luciferase reporter HEK 293T cells. (**d**) IFN-β promoter activity as function of PB2 mutation and mvRNA formation. In all panels, error bars indicate standard deviation. P-values were determined using ANOVA in Fig. 1a and c, or linear regression in Fig. 3d, and are indicated as * P ≤ 0.05, ** P ≤ 0.01, *** P ≤ 0.001, **** P≤ 0.0001.

The identification of mismatches and base-skipping during the generation of mvRNAs (see Fig. 2e) suggests that mvRNA production might be dependent on polymerase fidelity. To investigate this further, we introduced a V43I mutation, which has been shown to confer high-fidelity on an H5N1 influenza virus polymerase^22^, into the PB1 subunit of the BM18 polymerase (BM18^hf^). We found that mvRNA levels were significantly reduced in the presence of BM18^hf^, with a corresponding reduction in IFN-β promoter activity (Fig. 3a). Together, the observations in Fig. 2a, 3a and Fig. S3a suggest that dysregulation of viral RNA replication, e.g. by limiting NP availability, and replication by highly pathogenic avian influenza virus polymerases in mammalian cells generates mvRNAs by employing an error-prone copy-choice mechanism, such as proposed for recombination in positive-strand RNA viruses^23^.

Having observed a significant difference in mvRNA production by the WSN and BM18 polymerases, we next asked whether a particular subunit is the determinant of mvRNA production. Therefore, we replaced individual polymerase subunits of the BM18 polymerase with subunits of the WSN polymerase in the 246 nt RNA replication assay. We found that particularly replacement of the BM18 PB2 subunit with the WSN PB2 subunit eliminated the generation of mvRNAs (Fig. S4a). Interestingly, the BM18 influenza PB2 subunit has been linked to the enhancement of both the kinetics and the magnitude of the host response to viral infection, leading to the induction of strong inflammatory responses with increased cellular infiltration in the lungs of infected mice^24^. To identify PB2 amino acid residues involved in the formation of mvRNAs, we aligned the BM18, WSN, NT60 and FJ02 PB2 sequences and found four amino acid positions that distinguish the BM18 and FJ02 polymerases from the WSN and NT60 polymerases: 9 (D→N), 64 (M→T), 81 (T→M), and 661 (A→T) (Fig. S4a). Each of these amino acids has been implicated in avian to mammalian host adaptation^25^ and, interestingly, all three N-terminal PB2 adaptive amino acids map to the template exit channel of the RNA polymerase heterotrimer (Fig. 3b) ^6^. We generated single mutations N9D, T64M, M81T, and double mutations N9D+T64M and N9D+M81T in the PB2 subunit of the WSN polymerase and found that mutants N9D and M81T, and the double mutants N9D+T64M and N9D+M81T, significantly increased mvRNA formation (Fig. 3c) and IFN-β promoter activity (Fig. 3c and d). However, the levels of mvRNAs generated by these mutants did not reach the levels generated by the BM18 polymerase indicating that further amino acids are involved in determining the ability of a polymerase to produce mvRNAs. In line with our observations, WSN viruses that contain a PB2 N9D substitution or other PB2 mutations near the template exit channel have been reported to induce higher IFN-β expression than wild-type WSN^26,27^.

To address whether mvRNAs form during infection of mammalian cells, we infected A549 cells with WSN, the highly pathogenic avian strain A/Vietnam/1203/04 (H5N1) (abbreviated as VN04), and the VN04 virus with the PB1 V43I high-fidelity mutation (abbreviated as VN04^hf^). PAGE and deep sequencing analysis using RT-PCR universal primers (Fig. S2a) showed that infections with VN04 resulted in high levels of mvRNAs, while WSN infections produced only very low levels (Fig. 4a). Infections with VN04^hf^ resulted in significantly reduced mvRNA levels compared to the wild-type VN04 virus. These results demonstrate that mvRNAs are formed during influenza virus infection of lung epithelial cells and that polymerase fidelity is an important determinant of mvRNA formation (Fig. 4a).

**Figure 4.**
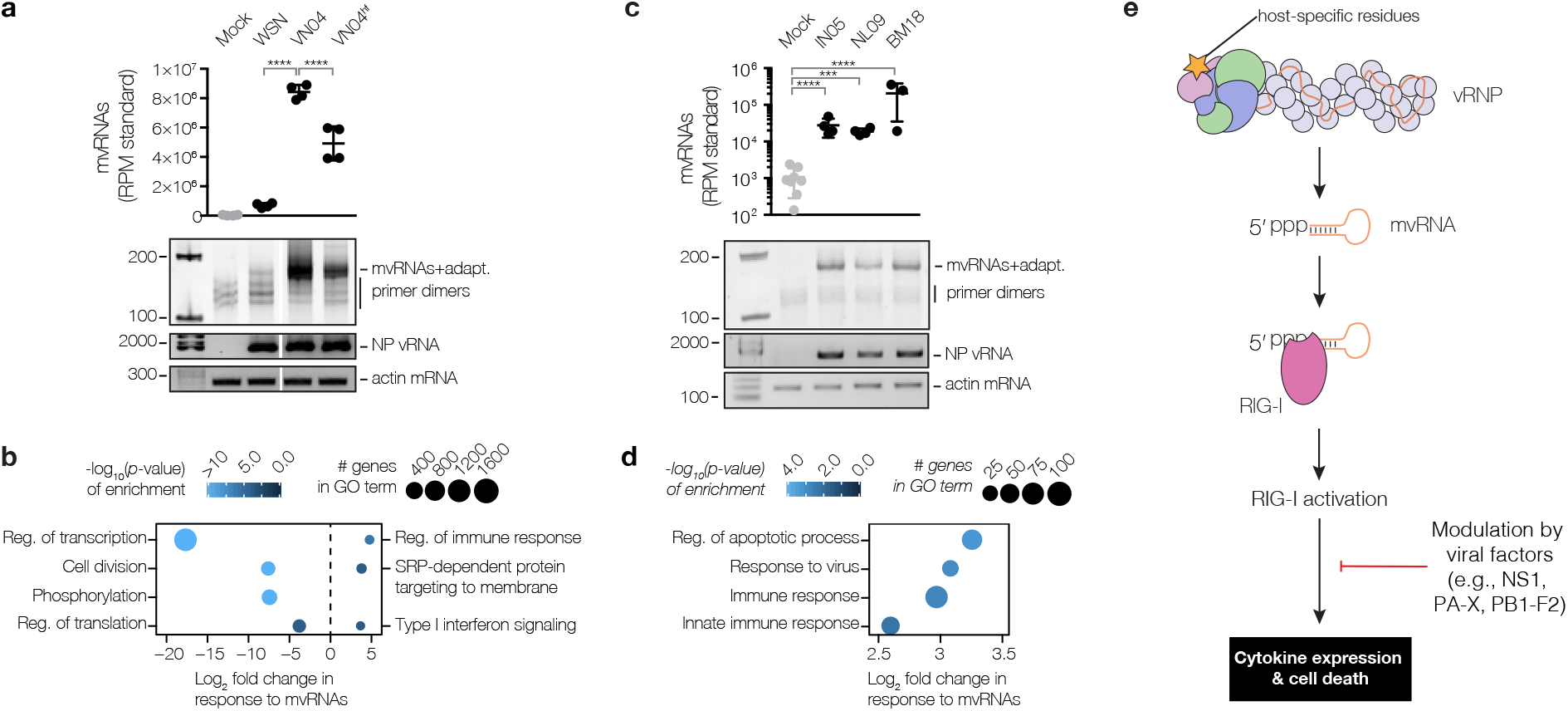
Levels of mvRNAs produced during infection correlate with innate immune responses. (**a**) Analysis of mvRNAs in A549 cells infected with WSN, VN04, VN04^hf^ using deep sequencing or PAGE. mvRNAs were amplified using universal primers containing adapters for sequencing (mvRNAs+adapt). mvRNA counts were normalised to mvRNA and mcRNA internal standards. (**b**) Analysis of mRNAseq of infected A549 cells showing GO terms down-regulated (left) and GO terms up-regulated (right) in VN04 infection as compared to VN04^hf^ in response to mvRNA levels. (**c**) Analysis of mvRNAs in lungs of ferrets one day after infection with IN05, NL09 or BM18 using deep sequencing or PAGE. (**d**) Analysis of tissue mRNAseq showing GO terms enriched as function of mvRNA levels in lungs of ferrets infected with IN05, NL09 and BM18 influenza viruses. (**e**) Model for the expression of cytokines in influenza virus infected cells. In all panels, error bars indicate standard deviation. P-values were determined using ANOVA and are indicated in Fig. 4a and c as in Fig. 3, and using a one-sample z-test (see Methods) for Gene Ontology analysis in Fig. 4b and d, and are indicated by blue shading.

To investigate whether there is a link between mvRNA production and virus-induced innate immune responses we performed RNAseq of cells infected with VN04 and VN04^hf^ viruses, which differ only at a single amino acid residue, and examined which genes were differentially expressed in response to mvRNA levels. Despite significantly different mvRNA levels produced by VN04 and VN04^hf^, viral mRNA levels were similar (Data S1), in agreement with previous findings that the V43I mutation has only a marginal effect on virus replication^22^. Gene Ontology (GO) analysis (Fig. 4b) showed that basic cellular functions (e.g. translation and transcription) were significantly compromised in VN04 infection relative to VN04^hf^, consistent with a greater level of cell death, which is known to exacerbate inflammation^28^. In addition, we observed that genes associated with innate immune responses showed a significant increase in expression in response to higher mvRNA levels (Data S1, Fig. 4b). Overall, these observations are indicative of a link between erroneous polymerase activity, mvRNA synthesis, and innate immune activation and the induction of cell death. Furthermore, as VN04 exhibited a 10-fold higher lethality compared to V04^hf^ in mice^22^, our data also suggest a link between mvRNA levels and virulence.

To address whether mvRNAs are also produced in infection of animal models, we investigated mvRNA generation in ferrets, which are regarded as the gold-standard model for influenza virus infection studies. Specifically, we analysed RNA samples from ferret lungs one and three days after infection with highly pathogenic avian A/Indonesia/5/2005 (H5N1) (abbreviated as IN05), 2009 swine-origin pandemic A/Netherlands/602/2009 (H1N1) (abbreviated as NL09) or the BM18 pandemic virus^29,30^. mvRNAs were present in all infected lung samples one day after infection, with mvRNA levels particularly high in the BM18 infected ferret lungs (Fig. 4c, Fig. S5). RNAseq and differential gene expression analysis, followed by GO analysis on the ferret lung samples taken one and three days post infection, showed an up-regulation of apoptosis and innate immune responses as function of mvRNA level, independently of viral titre or the day post infection (Fig. 4d).

In summary, we identify mvRNAs, a novel class of influenza virus RNAs, that act as the main agonists of the pathogen recognition receptor RIG-I during influenza virus infection (Fig. 4e). mvRNAs are produced as a result of aberrant replication of the viral RNA genome by the viral RNA polymerase. Polymerase fidelity and host-specific amino acids are determinants of the ability of the viral polymerase to produce mvRNAs, which are distinct from DI RNAs and full length viral RNA segments in that they can be efficiently replicated in the absence of NP and do not form canonical RNPs17. These features of mvRNAs are likely to be critical for their preferential recognition by RIG-I over DI RNAs and full length RNA segments. We further demonstrate that mvRNA production is linked to increased cytokine expression and cell death. Since high levels of mvRNAs lead to more innate immune activation and cell death, our observations strongly suggest that mvRNAs are a contributing factor to influenza virus virulence. We speculate that production of high levels of mvRNAs by the polymerases of the 1918 pandemic and highly pathogenic avian influenza viruses and the resulting increased innate immune activation contribute to the cytokine storm phenomenon underlying the high virulence of these viral strains. The effects of mvRNAs are likely modulated by viral factors, such as the immunomodulatory NS1 and PB1-F2 proteins^12^ (Fig. 4e). Further studies are required to assess mvRNA levels generated by various influenza virus strains, including seasonal strains, and their effect on virulence.

## Methods

### Ethics and biosafety

All work with highly pathogenic H5N1 viruses in A549 cells was conducted in the Biosafety Level-3 laboratory at the LKS Faculty of Medicine, The University of Hong Kong, under guidelines and ethics approved by the Committee on the Use of Live Animals in Teaching and Research (CULATR). Ferret experiments with IN05 and NL09 were described previously^29^ and conducted in the Biosafety Level-3 laboratory of the Erasmus Medical Centre in compliance with European guidelines (EU directive on animal testing 86/609/EEC) and Dutch legislation (Experiments on Animals Act, 1997), after approval by the independent animal experimentation ethical review committee of the Netherlands Vaccine Institute (permit number 200900201). Ferret experiments with BM18 were described previously^30^ and approved by Institutional Animal Care and Use Committee of Rocky Mountain Laboratories, National Institutes of Health, and conducted in an Association for Assessment and Accreditation of Laboratory Animal Care international-accredited facility according to the guidelines and basic principles in the United States Public Health Service Policy on Humane Care and Use of Laboratory Animals, and the Guide for the Care and Use of Laboratory Animals. Sample inactivation and shipment was performed according to standard operating procedures for the removal of specimens from high containment and approved by the Institutional Biosafety Committee.

### Plasmids

Plasmids expressing the three polymerase subunits and NP of influenza A/WSN/33 (H1N1) ^31^, A/Northern Territory/60/68 (H3N2)^32^, A/duck/Fujian/01/02 (H5N1) ^32^(all pcDNA3-based), and A/Brevig Mission/1/18 (H1N1) ^33^(pCAGGS-based) have been described. A PB2 E627K mutation was introduced into the A/duck/Fujian/01/02 (H5N1) PB2 subunit to enable the FJ02 polymerase to efficiently replicate vRNA in mammalian cells. Plasmids expressing mutant PB1a (D445A/D446A) ^34^, and mutant PB2 (N9D) ^26^, of influenza A/WSN/33 (H1N1) virus have been described previously. Full-length or internally truncated vRNAs were expressed from plasmids under the control of cellular RNA polymerase I promoter^35^. Luciferase reporter plasmid under the control of the IFN-β promoter (pIFΔ(−116)lucter), the β-galactosidase reporter plasmid (pJatLacZ) under the control of a constitutive promoter (β-gal), pcDNA-Myc-RIG-I expressing myc-tagged RIG-I, and pcDNA-myc-proEGF have been described previously^36,37^. To construct plasmids expressing mutant PB1, PB2 proteins and myc-RIG-I (myc-RIG-I mut; which contains the mutations K851A, K858A and K861A), the plasmids expressing wild-type proteins were subjected to site-directed mutagenesis using the primers listed in Table S1.

### Cells and antibodies

Human embryonic kidney HEK 293T cells were originally sourced from the ATCC, stored in the Dunn School cell bank at the University of Oxford, and mycoplasma tested, but not authenticated prior to our experiments. A549 cells were originally sourced from the ATCC and cultured at the University of Hong Kong. Cells were cultured in DMEM (Sigma-Aldrich) and 10% FCS. Western blots were performed using NP antibody GTX125989 (GeneTex), Myc antibody GTX115046 (GeneTex), RIG-I antibody GTX85488 (GeneTex), and PB2 antibody GTX125926 (GeneTex). Wild-type and RIG-I knockout HEK 293T cells expressing luciferase in response to the activation of the IFN-β promoter were described previously^38^.

### Statistical testing

In all figures, error bars indicate standard deviation with sample sizes as indicated in figures or figure legends. Evaluation of the statistical significance between group means was performed across all experiments according to the following criteria: (i) in the case where a comparison of a single variable was made between only two groups, an unpaired t-test was used; (ii) in the case of comparisons between three of more groups of measurements derived from a single independent variable (e.g. IFN-β induction as a function of RNA length), one-way ANOVA was used and P-values were corrected for multiple comparisons using either Dunnett’s test (when a single group was taken as a reference/control to which all other groups were compared) or the Bonferroni method (when specific pairs of groups were compared to one another); (iii) in the case of comparisons between three of more groups of measurements derived from two independent variables (e.g. IFN-β induction as a function of RNA length and RIG-I expression), two-way ANOVA was used and P-values were corrected for multiple testing using the Bonferroni method; (iv) in the case of comparisons between three of more groups of log-distributed data (e.g. viral titres), measured values were first log_10_ transformed and then compared using one-way ANOVA, with P-values corrected for multiple comparisons by controlling the false discovery rate (FDR) to be <0.05 using the two-stage step-up method of Benjamini, Kreiger, and Yekutieli. For the evaluation of the statistical significance of the relationship between two measured values (e.g. fold increase in IFN-β induction *vs.* mvRNA level), linear regression analysis was used, with the P-value indicating the probability of the null hypothesis (no linear relationship), and the goodness of fit reported as r^2^. Statistical testing related to differential gene expression analysis is detailed below, and was performed in R; all other statistical tests were performed using GraphPad Prism.

### RNP reconstitution assays and quantitative RNA analysis

RNP reconstitution assays were carried out in 24-well plates out as described previously^34,39^. Briefly, 0.25 µg of the plasmids pcDNA3-NP, pcDNA3-PB2, pcDNA3-PB1, pcDNA3-PA, and a pPOLI plasmid encoding full-length or truncated vRNA templates (for list of vRNA templates used see Table S2) were transfected into HEK 293T cells using Lipofectamine 2000 (Invitrogen) according to the manufacturer’s instructions. Twenty-four hours post transfection, RNA was extracted using TRI Reagent (Sigma-Aldrich) and dissolved in RNase free water. For quantitative primer extensions, reverse transcription was carried out using SuperScript III reverse transcriptase (Thermo Fisher Scientific) with ^32^P-labelled oligonucleotides complementary to vRNA-derived RNA species and ribosomal 5S rRNA (for primers see Table S3). cDNA synthesis was stopped with 10 µl loading dye (90% formamide, 10 mM EDTA, xylene cyanole, bromophenol blue) and ^32^P-labelled cDNAs generated with primer NP-were resolved by 12% denaturing PAGE (19:1 acrylamide/bisacrylamide, 1x TBE buffer, 7 M urea). ^32^P-labelled cDNAs generated with primer NP-2 were resolved by 20% denaturing PAGE. The radiolabelled signals were imaged using phosphorimaging on a FLA-5000 scanner (Fuji), and analysed using AIDA (RayTek) and Prism 7 (GraphPad). In all experiments, the apparent RNA levels were background corrected using the PB1 active site mutant (PB1a) signal and normalised to the 5S rRNA control. Statistical analysis of data from at least three independent experiments was carried out using ANOVA.

### RNP reconstitution assays and qualitative RNA analysis

For segment-specific qualitative RNA analysis by RT-PCR, RNA was treated with DNase (Promega) for 10 min according to the manufacturer’s instructions and reverse transcribed using SuperScript III and the PB2 primers listed in Table S3. cDNA was amplified using Q5 polymerase (NEB) and the primers listed in Table S3. PCR products were analysed on 1.5% agarose gels in 0.5x Tris-acetate-EDTA (TAE) buffer. For qualitative RT-PCR using universal primers, DNase treated RNA was reverse transcribed using the Lv3aa and Lv3ga primers listed in Table S3 and Superscript III at 37 ºC for 30 min. Second strands synthesis was performed with primer Lv5 and Q5 polymerase (NEB) at 47 ºC for 10 min, followed by a further extension at 72 ºC for 3 min. The primer excess in the reactions was removed by incubating the second strand reaction with 1 U of exonuclease VII (NEB) at 37 degrees Celsius for 1 h. Following inactivation of the exonuclease at 95 ºC for 10 min, the DNA was amplified using Q5 polymerase, and primers P5 and i7 for 25 cycles. PCR products were analysed by 6% PAGE.

### Luciferase-based interferon expression assays

For luciferase assays, RNP reconstitutions were performed in wild-type HEK 293T cells or HEK 293T cells engineered to express luciferase from the IFN-β promoter. RNP reconstitutions were performed in a 24-well format by transfecting 0.25 µg of the plasmids pcDNA3-NP, pcDNA3-PB2, pcDNA3-PB1, pcDNA3-PA, a pPOLI plasmid encoding full-length or truncated vRNA templates using lipofectamine2000 (Invitrogen). For RNP reconstitutions in wild-type HEK 293T cells, 100 ng of pIFΔ(−116)lucter and pJatLacZ were co-transfected with the polymerase expressing plasmids. Twenty-four hours post transfection, cells were harvested in PBS and resuspended in Reporter Lysis buffer (Promega). Luciferase activity was measured using a Luciferase Assay System (Promega) and a GloMax (Promega), and normalised using the β-galactosidase signal measured using ortho-Nitrophenyl-β-galactoside (ONPG) and a GloMax. The background was subtracted using signals obtained from cells transfected with an empty pcDNA3. Luciferase levels were corrected for viral RNA levels obtained with primer extensions and a ^32^P-labelled NP-2 primer (Table S3). For total RNA transfections, 100 ng of total RNA was transfected with 100 ng of pIFΔ(−116)lucter and pJatLacZ using Lipofectamine2000. Analysis of luciferase expression was performed as described above. Statistical analysis was carried out using ANOVA.

### Immunoprecipitations

For myc-RIG-I immunoprecipitations, 10 cm dishes with HEK 293T cell were transfected with 3 µg pcDNA3-NP, pcDNA3-PB2, pcDNA3-PB1, pcDNA3-PA, pcDNA-myc-RIG-I or pcDNA-myc-EGF, and a pPOLI plasmid encoding either a full-length or truncated vRNA template using Lipofectamine 2000. Twenty-four hours post transfection, the cells were harvested in cold PBS and lysed in 600 µl Tris lysis buffer (50 mM Tris-HCl, pH 8.0; 5% glycerol; 0.5% Igepal; 200 mM NaCl; 1 mM EDTA; 1 mM DTT; and 1x EDTA-free protease inhibitor (Roche)) on ice for 1 h. The lysates were cleared at 10,000 *g* for 5 min. Six µg of anti-myc antibody (Sigma-Aldrich) was added to 0.5 ml of cleared lysate and mixed at 4 ºC for 1.5 h. The lystate-antibody mix was bound to Dynabeads (Novex) at 4 ºC for 1.5 h, washed 3 times with IgG wash buffer (10 mM Tris-HCl pH 8.0; 150 mM NaCl; 0.1% Igepal; 1 mM PMSF; 1 mM EDTA), and finally analysed for bound RNA and protein. Statistical analysis of data from three independent experiments was carried out using ANOVA.

### ATPase assay

For wild-type and mutant myc-RIG-I purification, HEK 293T cell were transfected with 5 µg pcDNA-myc-RIG-I or pcRNA-myc-RIG-I mut using Lipofectamine 2000. Twenty-four hours post transfection, the cells were harvested in cold PBS and lysed in lysis buffer (50 mM Hepes, pH 8.0; 5% glycerol; 0.5% Igepal; 200 mM NaCl; 2 mM MgCl_2_; 10 mM CaCl_2_; 1 mM DTT; 1 U/ml Micrococcal Nuclease (Thermo Scientific); and 1x EDTA-free protease inhibitor) on ice for 1 h. Three µg of anti-myc antibody was next added per 0.5 ml of cleared lysate and mixed at 4 ºC for 1.5 h. The lystate-antibody mix was bound to Protein G Mag Sepharose Xtra beads (GE Healthcare) at 4 ºC for 1.5 h, washed 6 times with 20 column volumes of RIG-I wash buffer (50 mM Hepes, pH 8.0; 200 mM NaCl; 0.1% Igepal; 5% glycerol; 1 mM PMSF; 2 mM MgCl_2_) at 4 ºC for 10 min, and finally myc-RIG-I was eluted from beads in 1 column volume wash buffer containing 0.5 mg/ml c-myc peptide (Pierce) for 15 min at 4 ºC. Activity assays were performed in 50 mM Hepes pH 8.0, 150 mM NaCl, 2 mM MgCl_2_, 5 mM DTT, and 0.1 μM [γ-^32^P]ATP. [γ-^32^P]ATP and ^32^P_i_ were resolved using PEI-cellulose TLC plates (Sigma-Aldrich) in 0.4 M KH_2_PO_4_ pH 3.4.

### Cell and animal infections

HEK 293T cells were infected with influenza A/WSN/33 (H1N1) virus, free of DI RNAs, at a multiplicity of infection (MOI) of 5. RNA was extracted 5 h post infection and analysed using deep sequencing or qualitative RT-PCR. 6-wells containing A549 cells were infected with A/HK/68 (H3N2), A/OK/1992/05 (H3N2), A/Brisbane/59/2007 (H1N1), A/Vietnam/1203/04 (H5N1), or A/Vietnam/1203/04 (H5N1) with the V43I mutation with an MOI of 5. RNA was extracted 8 hours post infection and analysed using deep sequencing or qualitative RT-PCR. Ferret lung tissue was obtained from ferrets infected with A/Indonesia/5/2005 (H5N1) or A/Netherlands/602/2009 (H1N1) ^29^, or A/Brevig Mission/1/1918 (H1N1) ^30^. A single ferret (lung titre = 7.6×10^1^ log_10_TCID_50_/g) was excluded from analysis on the basis of its apparent lack of infection. Ferret RNA was isolated from lung tissue samples using Trizol (Invitrogen) and analysed using qualitative RT-PCRs and next generation mvRNA sequencing with universal primers and quantitative mRNA sequencing.

### Sequence alignment and structural modelling

PB2 amino acid sequences from influenza A viruses A/WSN/33 (H1N1), A/Brevig Mission/1/18 (H1N1), A/Northern Territory/60/68 (H3N2), and A/duck/Fujian/01/02 (H5N1) were aligned using Muscle 3.0 and visualised using ESPript^40^. The bat influenza A virus polymerase structure (PDB 4WSB) was visualised in Pymol 1.6.

### Next generation sequencing of mvRNAs using adapters

Total cell RNA from transfected or infected cells was isolated using Tri Reagent (Sigma) or Trizol (Invitrogen) according to the manufacturer’s instructions and fractionated into small (17-200 nt) and large (>200 nt) RNA fractions using an RNA Clean and Concentrator kit (Zymo Research). Next, the small RNA fraction was denatured at 70 ºC for 2 min and subsequently treated with 2 U of XRN-1 in NEB buffer 2 at 37 ºC for 15 min to deplete miRNAs. Next, XRN-1 was inactivated by adding 10 mM EDTA and incubating the reaction at 70 ºC for 10 min. Viral triphosphorylated RNAs were converted to monophosphorylated RNAs by adding 5 U of RNA 5’ Pyrophosphohydrolase (RppH) and 10 mM MgCl_2_ and incubating the reactions at 37 ºC for 15 min. RNA was purified using an RNA Clean and Concentrator kit and libraries for deep sequencing were prepared using the NEBNext Small RNA Library Prep Kit according to the manufacturer's instructions. To ensure accurate quantitation after PCR amplification, the concentration of each library was measured by qPCR on a StepOnePlus instrument (ABI) and the number of PCR cycles used to subsequently amplify the remaining library material was calibrated so as to ensure the PCR was in the early stage of exponential amplification and to not over-cycle the PCR reactions. Amplified sequencing libraries were purified on a 6% Novex TBE PAGE according to the manufacturer’s instructions to remove primer-dimers. Paired-end sequencing (2×75bp) on an Illumina HiSeq 4000 was carried out by the Oxford Genomics Centre, Wellcome Trust Centre for Human Genetics (Oxford, UK).

### Next generation sequencing of mvRNAs using universal primers

To spike viral RNA for quantitative sequencing, 0.2 µl of 100 pM spike RNA (Table S2) was added to 40 ng of the small RNA fraction (see above). The RNA mixture was next converted into cDNA using primers Lv3aa, Lv3ga and Lc3 and Superscript III (Invitrogen) at 37 ºC for 30 min. Second strand synthesis was performed using Q5 polymerase (NEB) and primers Lv5, Lc3a, and Lc3g at 47 ºC for 10 min, followed by a further extension at 72 ºC for 3 min. The excess of barcoded primers was removed by incubating the second strand reaction with 1 U of exonuclease VII (NEB) at 37 ºC for 1 h. The exonuclease was inactivated at 95 ºC for 10 min. Next, the DNA was amplified using Q5 polymerase, primer P5 and i7 index primers (Lexogen), and subsequently sequenced on a NextSeq 500 sequencer (Illumina).

### Preparation of reference genome files for deep sequencing of mvRNAs

Prior to mapping, a reference genome file was prepared from relevant viral reference sequences in Genbank (see above). For the analysis of sequencing libraries prepared using universal influenza virus primers, the 5′ and 3′ viral promoter sequences of each segment were modified to match the degenerate universal primer sequences used in sample preparation (see Table S3), and the sequences of the spiked-in mvRNA quantitation standards (see Table S2) were appended to the reference genome. For WSN, VN04, and VN04^hf^ viruses, deep-sequencing data of mRNA generated in A549 cell infections (above) was exploited to generate updated reference genome files: the *mpileup* and *consensus* commands in the *bcftools* software package^41^ were used following mapping of non-host mRNA reads to the relevant viral reference genome using *STAR* aligner^42^.

### Data processing pipeline for deep sequencing of mvRNAs using universal influenza virus primers

Raw sequencing reads were first trimmed to remove sequencing adaptor sequences and reads with quality scores less than 20 using the *cutadapt* software package^43^ and the 8-nt unique molecular identifier (UMI) at the start of each read were removed from the sequence and appended to the read ID line of the FASTQ file using the *extract* command from the *umi_tools* software package^44^. Sequencing reads were then mapped end-to-end to the appropriate viral reference genome using the *STAR* aligner^42^, and permitting sequencing reads to have long internal deletions (i.e. an mvRNA, interpreted as splicing by STAR) with at least 16 nt anchored on either side of the deletion (*--outSJfilterOverhangMin 16 16 16 16*). The default settings of *STAR* were modified so that no alignment scoring penalty was given for an internal deletion and no preference was given to internal deletions that overlapped particular sequence motifs (*--scoreGapNoncan 0 --scoreGapGCAG 0 --scoreGapATAC 0*), and to ensure accurate quantitation, only the top-scoring alignment was included in the outputted BAM file (*--outSAMmultNmax 1*), which was sorted and indexed using *samtools*^45^. An aligned read was counted as an mvRNAs if it was anchored to the viral reference genome at the 5′ end in vRNA sense and contained an internal deletion (called as a splice junction by *STAR*), and the total numbers of mvRNAs and spiked-in quantitation standards were tallied using the *idxstats* command of *samtools*. mvRNA levels relative to the quantitation were then reported as number of reads per million mapped (RPM) quantitation standard. To validate the qPCR protocol used to prevent over-cycling of sequencing libraries, the quantitation was then repeated following removal of PCR duplicates, exploiting the UMI appended to each read, using the *umi_tools dedup* command, and counting the various mvRNA species as unique using the *--spliced-is-unique* option. The identities of the individual mvRNAs were then extracted from the *SJ.out.tab* file generated by STAR.

### Data processing pipeline for deep sequencing of mvRNAs using adapter ligation

Raw sequencing reads were first trimmed using the *cutadapt* software package^43^ to remove RNA adapters, sequencing adapters, and reads with quality scores less than 20. Since adapter ligation captures both host-derived and virus-derived small RNA species, reads were first mapped end-to-end to the DASHR database of human small RNAs^46^ using the *STAR* aligner^42^, with spliced alignments disabled (*--alignIntronMax 1*). Non-human reads were outputted using the *--outReadsUnmapped Fastx* option, were then mapped to the appropriate viral reference genome to find mvRNAs as described above, and quantitated relative to the total number of viral reads (RPM viral) or host reads (RPM host).

### Quantitative mRNA sequencing and differential gene expression analysis

Libraries for gene expression analysis were prepared using a QuantSeq 3' mRNA-Seq Library Prep Kit FWD for Illumina (Lexogen) according to the manufacturer’s instructions and sequenced on a NextSeq 500 sequencer. mRNA reads were aligned to the reference genome (CRCh38, GRCm38, or MusPutFur1.0) using the STAR read aligner^42^, exploiting the built-in trimming functions to remove the first 12 bases corresponding the Lexogen random primer (*--clip5pNbases 12*) and any contaminating poly(A) tails in the sequencing reads (*--clip3pAdapterSeq AAAAAAAAAAAAAAAAAA*), as well as requiring a minimum match to the reference genome of 40 bp (*--outFilterMatchNmin 40*). Gene counts were generated using reference genome annotations (Gencode v26 for CRCh38, and Ensembl 90 for MusPutFur1.0) using the STAR command *--quantMode GeneCounts*. Differential gene expression analysis was then carried out using the DEseq2 package in R^47^ to identify genes that were up- or down-regulated as a function of mvRNA levels, independently of viral load or titre. Specifically, the likelihood ratio test (LRT) was used to compare a full model (in which gene expression varies as a function of both viral load or titre, and mvRNA levels) to a reduced model (in which changes in gene expression are fully explained by viral load or titre alone) using analysis of deviance (ANODEV) to generate a P-value for the log-fold-change of each gene, which were adjusted for multiple testing by controlling the false discovery rate (FDR) using Independent Hypothesis Testing^48^ and reported as q-values. mvRNA levels were determined by deep sequencing using universal influenza virus primers, as detailed above. Viral load or titre was determined by segment 6 qRT-PCR or by using previously published values^29,30^. Subsequent enrichment analysis of Gene Ontology terms specifically affected by mvRNA levels was carried out using Parametric Analysis of Gene Set Enrichment^49^ via the GAGE package in R, with data from the above genome annotations, accessed via the biomaRt package^50^. Significance of enrichment for GO terms was calculated using a one-sample z-test^49^ in GAGE, and P-values were adjusted for multiple testing using the Benjamini-Hochberg method and were reported as q-values.

## Acknowledgements

We greatly value our discussions with R. Sun and Y. Du, who independently found that mutations in the N-terminal region of PB2, near the template exit channel of the polymerase, stimulate interferon induction^27^. We thank G.G. Brownlee, M. Freeman and F. Vreede for plasmids, J. Rehwinkel and A. Mayer for HEK 293 RIG-I -/- cells, Y. Kawaoka for the A/Brevig Mission/1/1918 (H1N1) virus, and A. Osterhaus and T. Kuiken for A/Indonesia/5/2005 (H5N1) or A/Netherlands/602/2009 (H1N1) infected ferret tissue samples. We thank Ian Sudbery for adding spliced read functionality to the umi_tools package. We thank the High-Throughput Genomics Group at the Wellcome Trust Centre for Human Genetics (funded by Wellcome Trust grant 090532/Z/09/Z) for the generation of adapter-ligated mvRNA sequencing data. This work was supported by Wellcome Trust grant 098721/Z/12/Z, joint Wellcome Trust and Royal Society grant 206579/Z/17/Z, and a Netherlands Organization for Scientific Research (NWO) grant 825.11.029 to A.J.W.t.V; EPA Cephalosporin Junior Research Fellowship (to D.L.V.B.); support by the Intramural Research Program of NIAID, NIH, to E.d.W.; Research Grants Council of the Hong Kong Special Administrative Region, China, Project No. T11-705/14N to L.L.M.P; and Medical Research Council (MRC) programme grants MR/K000241/1 and MR/R009945/1 to E.F. and studentship to J.C.L.

## Author Contributions

J.C.L., E.F., J.S. and A.J.W.t.V showed that subgenomic influenza virus RNAs stimulate IFN-β production and are bound by RIG-I. A.J.W.t.V. and J.C.L. found that mvRNAs are produced by influenza virus polymerases. D.L.V.B. designed sequencing strategies. D.L.V.B. and A.J.W.t.V. performed deep sequencing experiments and analyses. M.J.K., M.J.O.-M., H.F., and R.E.R. contributed reagents and protocols. E.d.W., D.v.R., and J.Y.S. provided ferret lung tissues. R.L.Y.F., H.Y. and L.L.M.P. performed A549 infections. A.J.W.t.V., D.L.V.B., J.C.L., and E.F. analysed data. A.J.W.t.V., J.C.L., D.L.V.B., and E.F. wrote the manuscript with input from co-authors.

## Materials and correspondence

All sequencing data have been deposited in the NCBI Sequence Read Archive (SRA) under accession number SUB3758924. Correspondence and requests for materials should be addressed to E.F. or A.J.W.t.V.

